# Pervasive short hairpin folding in non-encapsidated RNA viruses

**DOI:** 10.64898/2026.10.23.761338

**Authors:** Rob J. Dekker, Wim C. de Leeuw, Marina F. van Olst, Wim A. Ensink, Selina M. van Leeuwen, Timo M. Breit, Martijs J. Jonker

**Affiliations:** RNA Biology research group, Swammerdam Institute for Life Sciences, Faculty of Science, University of Amsterdam, Amsterdam, the Netherlands

**Keywords:** RNA viruses, RNA secondary structure, hairpin, stem-loop, ormycovirus, narnavirus

## Abstract

Genome-wide RNA secondary structure can influence viral replication, stability, translation, and interactions with host defenses, yet its distribution and significance across RNA viruses remain poorly understood. We developed the pervasive hairpin (peHaPi) metric to quantify densely distributed stem-loop structures and applied it to 817,234 GenBank virus sequences. This revealed a distinctive RNA architecture characterized by numerous short, highly uniform hairpins spanning most of the genome. The structure was strongly enriched in capsidless RNA viruses, encompassing nearly all ormycoviruses and a substantial fraction of narnaviruses, although it also occurred in several other RNA and DNA virus groups. Comparative analyses showed that increasing peHaPi scores appeared associated with more stable and constrained predicted RNA structures and altered codon base-pairing patterns, indicating selective constraints imposed by pervasive folding. Relationships between peHaPi and RdRp sequence similarity differed among virus groups, consistent with independent evolutionary origins of this structural phenotype. Analysis of virus-derived small RNAs provided no evidence that peHaPi(+) viruses elicit a reduced siRNA response, arguing against siRNA evasion as its primary function. Finally, combining peHaPi classification with DNA-read subtraction identified candidate RNA virus contigs lacking detectable protein homology or recognizable RdRp signatures across multiple independent datasets. These findings reveal pervasive short-hairpin organization as a novel feature of predominantly capsidless viruses and establish genome-wide RNA structure as a complementary signature for discovering highly divergent ormyco-and narnaviruses.

**Repositories:** The raw sequence reads have been deposited in the NCBI Sequence Read Archive under BioProject accession number <u>PRJNA1549704</u>.

## Introduction

Beyond its protein-coding role, RNA performs many other functions, including acting as a structural scaffold, regulating gene expression, carrying genetic information, and functioning as a catalytic molecule. [1]. All these capabilities rely heavily on its primary sequence as well as its secondary and tertiary structures, with functionality emerging from the interplay of local and long-range cis interactions [2]. The folding propensity of RNA stems from a diverse vocabulary of base-pairing modes, encompassing both canonical Watson–Crick pairs and a wide spectrum of non-canonical interactions [3, 4]. At the level of secondary structure, RNA forms stems and hairpins (stem-loops), internal loops, bulges, multibranch junctions, and pseudoknots [5–7]. Tertiary structure builds on these motifs through coaxial stacking, kissing-loop interactions, A-minor contacts, and diverse long-range tertiary bridges, which are stabilized by RNA-binding proteins [7–9]. Together, these interactions produce compact, information-dense structures whose conformations enable precise, sequence-driven functions. It is important to recognize that most RNA molecules exist as dynamic ensembles rather than fixed structures, with their conformations varying according to ionic conditions, temperature, crowding, and interactions with other macromolecules [10].

Few examples better illustrate the link between RNA structure and function than ribosomal RNAs. Their conserved secondary motifs fold into intricate 3D scaffolds via base triples, coaxial stacking, and metal-mediated bridges. These architectures stabilize ribosomal organization and precisely position catalytic residues within the peptidyl-transferase center, enabling peptide-bond formation during translation [11, 12]. Likewise, tRNAs fold from a cloverleaf secondary structure into a conserved L-shaped tertiary form, aligning the anticodon stem-loop with the 30S decoding center and the 3′ CCA end with the 50S peptidyl-transferase center [11, 13, 14].

Viral RNAs vividly illustrate structure-function coupling. For example, ∼70% of the hepatitis delta virus genome folds into a single unbranched rod that embeds self-cleaving ribozymes and exposes structures that regulate antigen expression [15, 16]. Similarly, coronavirus harbours frameshifting pseudoknots, 3′-UTR xrRNAs, retroviral TAR/DIS/Ψ hairpins, influenza panhandles, and picornavirus/hepacivirus IRES folds, which directly control replication, translation, and immune evasion [17–23].

Among RNA viruses, single-stranded RNA (ssRNA) bacteriophages exemplify how genome structure directs function [24]. Lacking packaging motors or pressurized capsids, these phages rely on structured genomic RNA rich in stem-loops that recruit maturation and coat proteins for assembly, with a principal operator hairpin guiding encapsidation [11, 25]. Additional dispersed stem-loops act as secondary packaging signals, aligning with a co-replicational assembly model integrating RNA folding, translation, replication, and encapsidation [26]. In the archetypal ssRNA phage MS2, coat proteins bind operator hairpins to repress replicase translation and tag RNA for packaging, while multiple binding sites coordinate capsid formation. Phage Qβ uses similar mechanisms with distinct replicase biology [27, 28]. Overall, compact ssRNA phage genomes encode many structural hairpins that integrate translation, replication, and virion assembly.

Virology distinguishes enveloped viruses (with a protein capsid plus lipid envelope) from non-enveloped (protein capsid only) and capsidless ones that do not produce a protein capsid or virion at all. Many encapsidated ssRNA viruses use distributed packaging signals that act cooperatively through multiple stem-loops and guide capsid assembly across diverse hosts [26, 29, 30]. Others, like members of the families *Narnaviridae, Mitoviridae*, and *Endornaviridae*, as well as ormycoviruses (here referring to viruses currently classified within the class *Orpoviricetes*) lack capsids entirely, replicating intracellularly as minimalist RNA replicons [31]. Some can even be transiently packaged by unrelated capsids, showing that RNA-capsid recognition is modular [32]. Even without encapsidation, structured cis-elements still govern replication, translation, and life-cycle switching [33, 34]. Recent research indicates that the single-stranded RNA genomes of narnaviruses exhibit extensive folding, suggesting that compact structural elements may be both widespread and functional even in capsidless systems where RNA-folding-based packaging signals are no longer necessary [35].

Despite all these insights it remains unclear whether capsidless RNA viruses might exploit their RNA folding structure to compensate for the absence of a protective capsid. In this study, we address this question by systematically analysing the RNA folding structure of capsidless viruses with a particular focus on narna(-like) viruses and ormyco(-like) viruses. In many of these capsidless viral RNAs, we discovered a characteristic, pervasive folding pattern in which short hairpins effectively cover the entire viral genome. By characterizing this genome-wide RNA folding in detail, we aim to elucidate how such structural organization may function as an alternative strategy to mitigate the lack of a canonical capsid.

Despite these insights, we lack a systematic, comparative view of short hairpin usage in (capsidless) RNA virus genomes. Do capsidless replicons, which do not need to present coat-protein binding motifs, nonetheless deploy short stem-loops broadly across their genomes? Here we address this knowledge gap by surveying RNA virus families for short stem-loops, integrating publicly available RNA virus genomes with RNA structure prediction and comparative sequence, codon-pairing, and smallRNA analyses. Our results support the view that compact stem-loops are ubiquitous in capsidless RNA virus genomes, point to potential roles in genome organization and intracellular RNA maintenance, and establish pervasive RNA folding as a useful signature for identifying highly divergent RNA viruses.

## Methods

### Biological samples and sequencing

Leaf samples were collected from 48 individual plants at Hortus Botanicus Leiden, the Netherlands, on 22 November 2024. Samples were immediately flash-frozen in liquid nitrogen and ground to a fine powder. Genomic DNA was isolated using the PureLink Genomic DNA Mini Kit (Thermo Fisher Scientific) according to the manufacturer’s instructions, while RNA was extracted from the same powdered material as described previously [36]. Library preparation and sequencing for RNA-seq, small RNA-seq, and DNA-seq were performed as described previously [36, 37]. Small RNA libraries were sequenced as 1 × 75-bp reads on an Illumina NextSeq 550, whereas RNA-seq libraries were sequenced as 2 × 150-bp reads on an Illumina NovaSeq X Plus.

### Read assembly, genomic DNA subtraction, and protein-homology search

Paired-end RNA-seq and DNA-seq reads were adapter-trimmed using Trimmomatic v0.39 [38] with `ILLUMINACLIP:2:30:10` and `MINLEN:36`. Trimmed paired RNA-seq reads were assembled separately for each sample using SPAdes genome assembler v3.15.5 [39] in RNA-virus mode (`--rnaviral`); for the Rhizoctonia and Erysiphaceae datasets, `--only-assembler` was additionally specified. Contigs shorter than 500 nt were removed prior to further analysis. Sequencing depth, SRA accessions for previously published datasets, and genomic references used for subtraction are summarized in Supplemental Table ST1.

To enrich for RNA virus-derived sequences, assembled contigs were screened for evidence of genomic DNA origin. For the *Rhizoctonia* and Erysiphaceae datasets, RNA contigs were aligned to the corresponding genomic reference(s) listed in Supplemental Table ST1 using minimap2 v2.22-r1105 [40] with `-x asm5 --secondary=no`. Contigs with an alignment containing more than 100 matching nucleotides were considered genome-supported and excluded. For the *Phytophthora* dataset, contigs were searched against the corresponding reference genome (Supplemental Table ST1) using BLASTN v2.13.0+ [41] with `-evalue 1e-20 - max_target_seqs 5`, and contigs with more than 50% query coverage were excluded.

For the botanic garden and seaweed datasets, matched DNA-seq reads were instead mapped directly to the corresponding RNA-seq contigs using Bowtie2 v2.4.1 [42] with the `--very-sensitive` preset. Both paired reads and reads rendered unpaired during trimming were included. Unmapped reads were discarded, and nucleotide coverage was determined using `samtools depth -aa` in SAMtools v1.20 [43]. Contigs were classified as DNA(−) only when no nucleotide was covered by a mapped DNA read.

Low-complexity regions were identified using dustmasker v1.0.0, distributed with BLAST v2.13.0 [44]. For the *Rhizoctonia*, Erysiphaceae, and *Phytophthora* datasets, retained candidates were at least 1,000 nt long and contained no more than 60% low-complexity sequence. For the botanic garden and seaweed datasets, retained candidates were longer than 1,000 nt and contained less than 90% low-complexity sequence.

Protein-coding sequences were predicted from candidate contigs using EMBOSS getorf v6.6.0.0 with a minimum ORF length of 150 nt and `-find 0`. Predicted proteins were searched against a locally stored clustered NCBI nr protein database (downloaded from NCBI on 08/07/2025) using DIAMOND blastp v2.1.10, retaining the single highest-ranking target per ORF (`--max-target-seqs 1`) with an E-value threshold of 1 × 10⁻⁵.

### RNA secondary-structure prediction and peHaPi scoring

RNA secondary structures were predicted using RNAfold from ViennaRNA v2.7.2 [45] with default folding parameters and `-p --noPS --noDP` options to enable additional thermodynamic ensemble calculations. For sequences longer than 5,000 nt, only the first 5,000 nt were analyzed, whereas shorter sequences were folded over their full length. The resulting minimum-free-energy dot-bracket structures were analyzed using custom R functions `TrimHairpin()`, `HairpinScore()`, and `LoopynessValue()` (<u>Supplemental Document SD1</u>) to quantify pervasive hairpin formation in R version 4.3.2 [46]. Briefly, simple hairpins were identified as structures containing a terminal loop of 3–9 unpaired nucleotides. Candidate hairpins were truncated at stretches of four or more consecutive unpaired nucleotides within the stem and balanced according to a specified maximum stem length, defined by the number of paired nucleotides on each side of the hairpin. The peHaPi score was calculated as the fraction of the analyzed sequence encompassed by the resulting hairpins (Supplemental Figure SF1). The maximum permitted stem length is indicated by the metric name; for example, peHaPi12 restricts each identified hairpin to a maximum of 12 paired nucleotides per stem. In addition, the standard deviation of total hairpin length in nucleotides was calculated for each sequence as a measure of within-sequence hairpin-size uniformity.

Hairpin density was calculated as the number of detected hairpins divided by the analyzed sequence length and expressed per 100 nt, using the custom function `HairpinScore()`. The number of unpaired nucleotides outside hairpins was calculated from the RNAfold dot-bracket structure by subtracting unpaired nucleotides contained within identified hairpins from the total number of unpaired nucleotides and normalizing the remainder by the number of detected hairpins using the custom function `NonHairPinUnpaired()` (<u>Supplemental Document</u> <u>SD1</u>).

### Optimization of peHaPi parameters by ROC analysis

The maximum permitted hairpin stem length used for peHaPi calculation was optimized by receiver operating characteristic (ROC) analysis using the R package pROC v1.19.1 [47]. For each tested maximum stem length, peHaPi scores were recalculated directly from the RNAfold dot-bracket structures using the method described above (<u>Supplemental Document SD1</u>). Maximum stem lengths from 3 to 50 nt were tested in 1-nt increments. To optimize discrimination of ssRNA phages, all 6,336 *Leviviricetes* sequences were assigned to the positive class and compared with a random sample of 10,000 non-*Leviviricetes* viral sequences assigned to the negative class. For each stem-length setting, an ROC curve was generated using `pROC::roc()`, the area under the ROC curve (AUC) was calculated using `pROC::auc()`, and the operating threshold was obtained using `pROC::coords(…, “best”, best.method = “youden”)`. This threshold maximizes Youden’s J statistic, defined as sensitivity + specificity − 1. The stem-length parameter yielding the highest AUC was selected, resulting in a maximum stem length of 29 nt (peHaPi29; AUC = 0.972) and an optimal peHaPi29 threshold of 0.514.

Discrimination between ormycoviruses and ssRNA phages was subsequently evaluated using the complete sets of 266 ormycovirus and 6,336 *Leviviricetes* sequences. ROC analyses were performed separately for peHaPi12 and the standard deviation of hairpin length. For peHaPi12, the Youden-optimized operating threshold was 0.573 (AUC = 0.986; sensitivity = 0.959; specificity = 0.942). The corresponding optimal threshold for hairpin-length standard deviation was 13.94 nt. These two criteria were subsequently combined to define peHaPi(+) sequences as peHaPi12 > 0.573 and hairpin-length standard deviation <13.94 nt.

### Minimum free energy and ensemble diversity analysis

Minimum free energy (MFE) and ensemble diversity were obtained from RNAfold predictions and normalized by the analyzed sequence length. Analyses included GenBank viral sequences ≥1,000 nt classified as ssRNA phages, narnaviruses, or ormycoviruses. To visualize their relationships with peHaPi12, sequences were ordered by peHaPi12 score separately within each virus class and divided into consecutive equal-frequency bins of 10 sequences. For each bin, the median, 25th and 75th percentiles, and 10th and 90th percentiles of MFE per nucleotide or ensemble diversity per nucleotide were calculated. To facilitate visualization, the binned median and quantile curves were smoothed using cubic smoothing splines (R function `smooth.spline`, `spar = 0.83`).

### Codon base-pairing phase analysis

Codon base-pairing phases were determined using GenBank CDS annotations of all narna- and ormycoviruses (downloaded 29 May 2026), together with their previously calculated RNAfold dot-bracket structures. When the RNAfold structure was shorter than the GenBank sequence, as occurred for sequences analyzed only over their first 5,000 nt, CDS analysis was restricted to the corresponding 5′ sequence region. RNAfold structures longer than the GenBank sequence were rejected.

For each CDS, coding positions were reconstructed in coding orientation from the GenBank location annotation, including complement and multi-segment locations, and the `/codon_start` qualifier was used to establish the reading frame. Positions outside the analyzed RNAfold region were discarded. After application of the `/codon_start` offset, only complete codons within uninterrupted CDS regions covered by the RNAfold structure were analyzed. Incomplete terminal codon fragments were excluded. Multiple and overlapping CDSs were analyzed independently, while individual nucleotide positions could retain membership of more than one overlapping CDS.

RNAfold base pairs were mapped to codon positions 1, 2, and 3 of the annotated CDSs. Only base pairs for which both nucleotides occurred within annotated coding regions contributed to the codon-phase analysis. The nine possible codon-position pairings were grouped into three phase classes: *c1* comprised 1:1, 2:3, and 3:2 pairings; *c2* comprised 1:3, 2:2, and 3:1 pairings; and *c3* comprised 1:2, 2:1, and 3:3 pairings. Pairing between different CDSs was permitted. For each CDS, a codon was counted once within a phase class when at least one of its nucleotides participated in a base pair assigned to that class, irrespective of the number of such base pairs. Because different nucleotides within the same codon could participate in different pairing phases, a codon could contribute to more than one phase class. Phase frequencies were expressed relative to the total number of complete codons retained for that CDS.

For visualization, narnavirus and ormycovirus sequences were ordered by peHaPi12 score and divided into consecutive equal-frequency bins of 20 observations. For each phase class, the bin-wise median and 25th–75^th^ percentile range were calculated. Bin-wise medians and interquartile ranges were first smoothed using a centered 7-bin moving average and subsequently interpolated using cubic smoothing splines (`smooth.spline, spar = 0.8`) evaluated at 500 evenly spaced points across the peHaPi12 range.

### RdRp palmprint extraction, sequence clustering, and multidimensional scaling

RdRp palmprint sequences were identified from the GenBank virus dataset using the following workflow: ORFs were translated with EMBOSS getorf v6.6.0.0 using a minimum ORF size of 150 nt and `-find 0`, which reports translated regions between stop codons. The resulting proteins were screened with PalmScan2 v2.0 [48] using its PSSM-based search `(-search_pssms`). When multiple PalmScan hits occurred in one viral genome, only the highest-scoring hit was retained. For each retained RdRp candidate, the palmprint was extracted using the reported A, B, and C motif sequences to define the boundaries within the predicted ORF. Palmprint sequences ≤90 aa were excluded.

To reduce redundancy, complete viral nucleotide sequences were clustered with MMseqs2 release 18-8cc5c [49] using `easy-cluster`, `16 threads`, and `-c 0.9`, with all other parameters at default. One MMseqs2-assigned representative genome per cluster was retained, provided it contained a palmprint >90 aa. Narnavirus, ormycovirus, and ssRNA-phage representatives were retained irrespective of peHaPi status, whereas “Other” viruses were retained only when classified as peHaPi(+), yielding 4,979 representatives.

To reduce overrepresentation of peHaPi(−) narnavirus and ssRNA-phage sequences, these groups were independently subsampled in R using `sample()` until peHaPi(−) and peHaPi(+) counts were equal within each virus class. This removed 737 peHaPi(−) narnavirus and 3,705 peHaPi(−) ssRNA-phage representatives, resulting in 537 sequences. Ormycovirus and “Other” representatives were not subsampled.

The 537 palmprint amino-acid sequences were aligned in R v4.3.2 using ClustalW [50] as implemented in msa v1.34.0 [51] with default parameters. The alignment was converted using ape v5.8.1 [52], and pairwise identity-based amino-acid distances were calculated with `dist.alignment(…, “identity”)` from the R package seqinr [53]. Classical multidimensional scaling was then performed with `cmdscale()` using two dimensions (k = 2). Biostrings v2.70.3 [54] was used for amino-acid sequence handling.

### RNA-seq and smallRNA-seq read mapping

Small-RNA reads were adapter-trimmed as described above and filtered to a minimum length of 19 nt. RNA-seq and small-RNA-seq reads were mapped to the 1,802 GenBank peHaPi(+) virus sequences and, where applicable, to *de novo* assembled viral contigs using Bowtie2 v2.4.1 [42] with default mapping parameters. A read was considered indicative of presence when its cumulative abundance across all tested samples was at least 1 for RNA-seq data and 10 for smallRNA-seq data.

### Hairpin-loop sequence analysis

Analysis of single nucleotide, triplet, and tetramer frequencies within the terminal hairpin loop regions from peHaPi(+) and peHaPi(‒) narna(-like) viruses was performed using custom R functions THL_Sequences(), KmerTable(), and THL_KmerTable() (Supplemental Document SD1).

### RdRp detection

In addition to PalmScan2, RdRp signatures were assessed independently using HMMER hmmsearch v3.3.2 [55] against the NeoRdRp v2.1 HMM database (Sakaguchi et al.), using an E-value threshold of 0.1 for the latter.

## Results and discussion

### Distinguishing genome-scale ordered RNA folding structures in ssRNA phages

Single-stranded RNA (ssRNA) phages are distinctive in that their RNA sequences are stably folded across their entire genome into a so-called genome-scale ordered RNA structure (GORS). This feature is exemplified by the well-studied ssRNA phages MS2 and Qβ (Figure 1A and B). To investigate whether similarly pervasive genome-wide folding occurs in other viral genomes, we developed a simple metric to quantify predicted stem-loop (hairpin) structures in any GenBank viral sequence of at least 1,000 nucleotides (n = 817,234; Supplemental Table ST2). This metric uses RNAfold [45] predictions for up to the first 5,000 nucleotides of each sequence and is reported as the pervasive hairpin (peHaPi) score, defined as the percentage of nucleotides predicted to form hairpin structures within that sequence (Supplemental Figure SF1).

**Figure 1.**
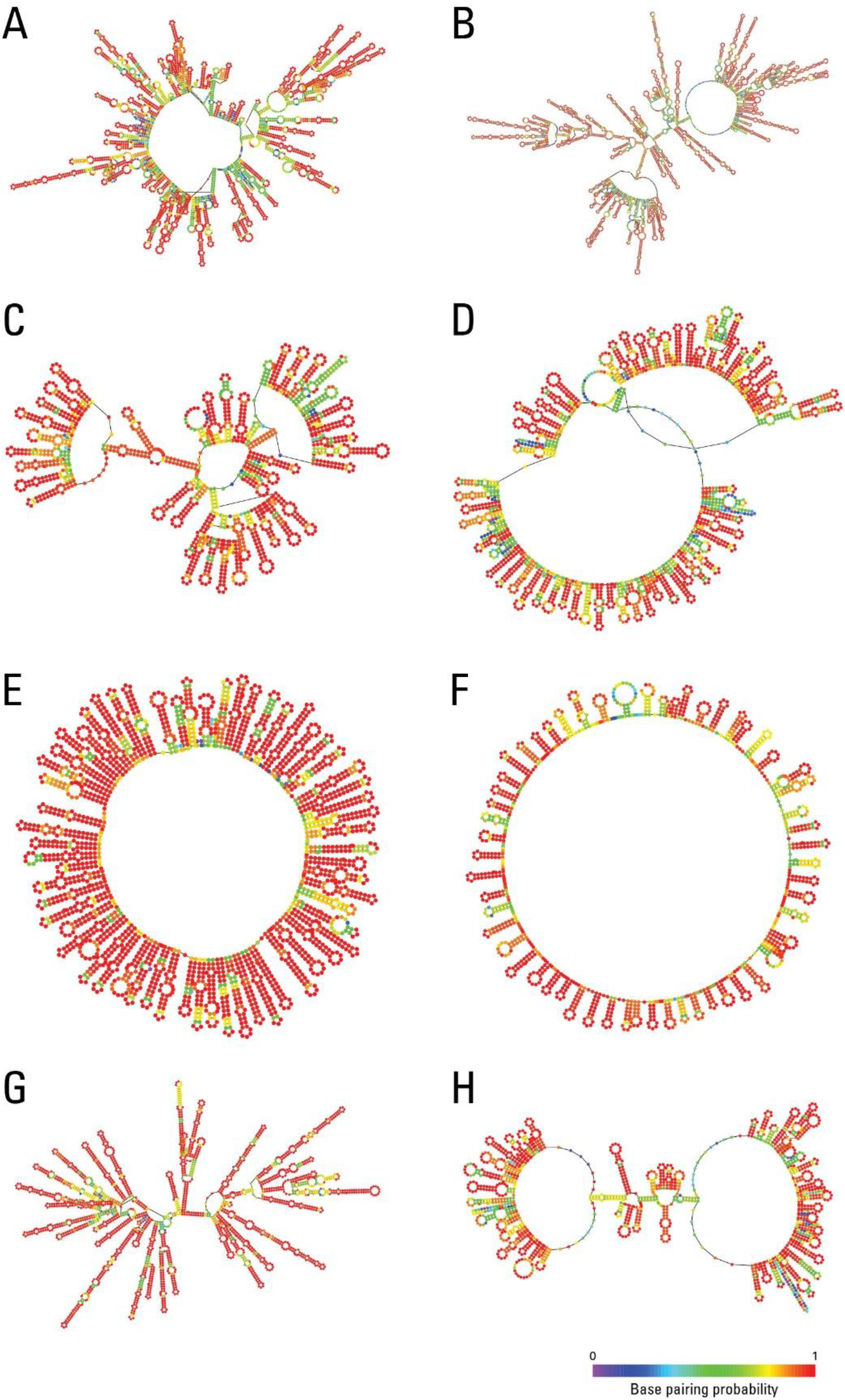
Representative examples of virus and phage genomes showing extensive RNA folding. Centroid secondary structures generated by RNAfold are shown for (A) ssRNA phage MS2 (NC_001417.2), (B) ssRNA phage Qβ (NC_001890.1), (C) phytophthora cinnamomi ormycovirus (PP891715.1), (D) Alaria esculenta RNA virus 1 (PP793780.1), (E) Sjack associated narna-like virus 18 (OQ987746.1), (F) grapevine wood holobiome associated narna-like virus 12 (PV021992.1), (G) Saccharomyces 20S RNA narnavirus (OK412912.1), and (H) cauliflower mosaic DNA virus (DQ870914.1). The color bar encodes base-pairing probabilities (0 – 1).

An important adjustable parameter of the peHaPi metric is the maximum number of paired nucleotides permitted in a hairpin stem. To optimize the ability of our metric to specifically identify ssRNA phage sequences based on their RNA folding patterns, we conducted a parameter sweep across maximum hairpin lengths ranging from 10 to 50 nucleotides (Supplemental Table ST3). Receiver operating characteristic area under the curve (ROC–AUC) values were computed for a benchmark dataset comprising 6,336 *Leviviricetes* ssRNA phage sequences and 810,898 non-*Leviviricetes* viral sequences obtained from GenBank. Using a maximum hairpin length of 29 nucleotides (peHaPi29) yielded the highest AUC (0.972). At the operating threshold that maximizes Youden’s J statistic (TPR-FPR), corresponding to a cut-off of 51.4%, we obtained a sensitivity of 0.948 and a specificity of 0.933 (Supplemental Table ST3). To illustrate the characteristic RNA folding signature of ssRNA phages, we plotted density distributions of peHaPi29 scores for both groups. ssRNA phage sequences displayed a pronounced peak at 65%, clearly separated from other viral sequences, which showed a peak at 38% (Figure 2A). Together, these results demonstrate that the peHaPi metric reliably identifies sequences exhibiting pervasive genome-wide folding.

**Figure 2.**
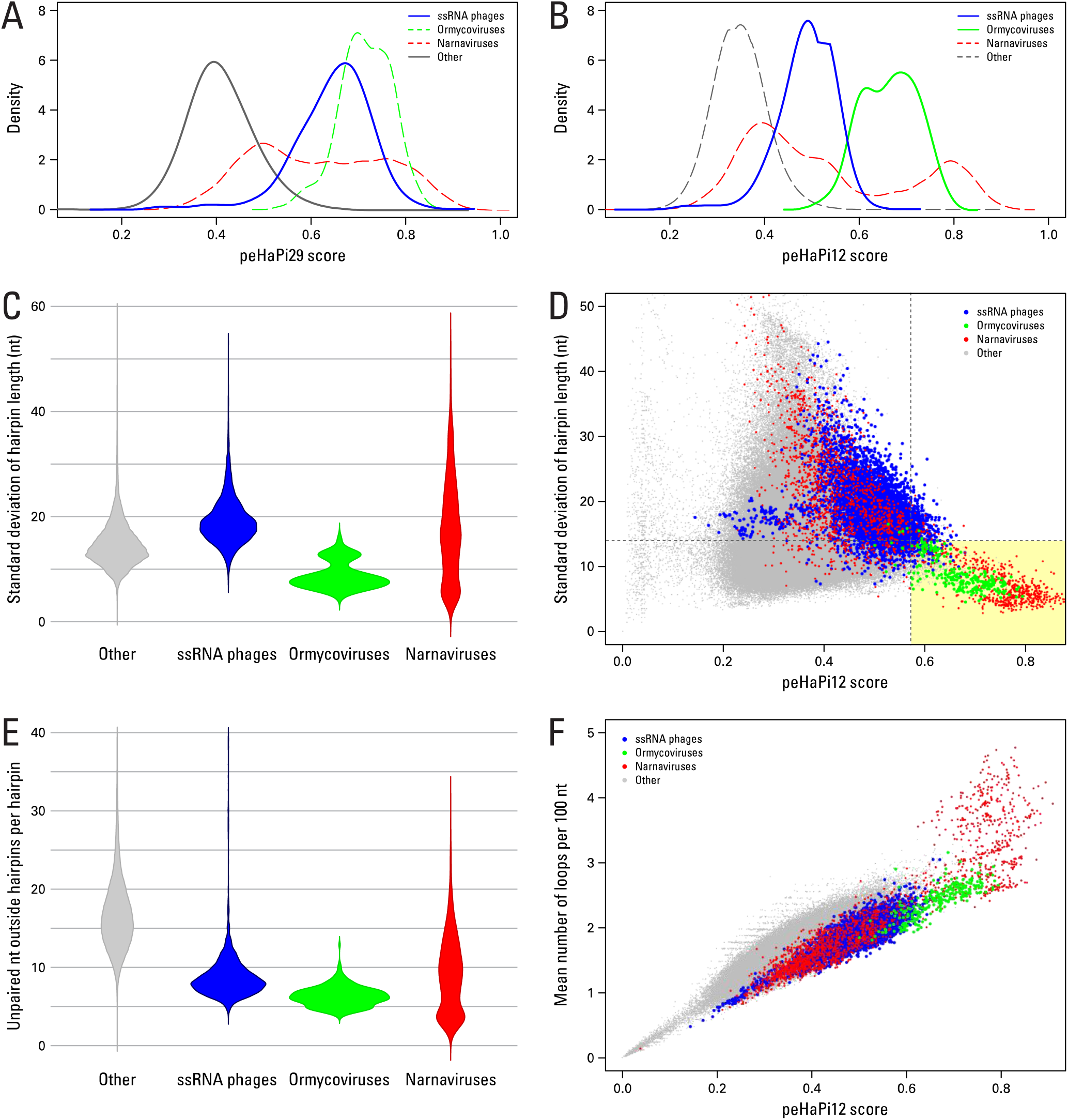
Distinctive peHaPi and RNA secondary structure features of ssRNA phages compared with other RNA viruses. peHaPi29 (A) and peHaPi12 scores (B-F) were calculated for all 817,234 GenBank viral sequences of at least 1,000 nucleotides (<u>Supplemental Table ST2</u>). Viruses were classified as ssRNA phages (blue), ormycoviruses (green), narnaviruses (red), and other viruses (gray). (A) Kernel density distributions of peHaPi29 scores (maximum hairpin length 29 nt). (B) Kernel density distributions of peHaPi12 scores (maximum hairpin length 12 nt). (C) Violin plots showing the distribution of the standard deviation of hairpin length. (D) Relationship between the peHaPi12 score and the standard deviation of hairpin length across virus genomes, with each point representing an individual viral genome. The vertical dashed line denotes the peHaPi12 optimal discrimination threshold (0.573), while the horizontal dashed line indicates the optimal cutoff for hairpin length standard deviation that maximizes the true positive rate/false positive rate trade-off (13.94 nt). ssRNA phage outliers (blue) with peHaPi12 scores below approximately 0.4 are attributable to large sequence gaps, represented as stretches of Ns in the corresponding GenBank genome records. (E) Violin plots showing the distribution of the number of unpaired nucleotides outside hairpins normalized over total number of hairpins per genome. (F) Relationship between peHaPi12 score and the mean number of loops per 100 nt.

### Discovery of a novel pervasive RNA hairpin structural type in non-phage viruses

Application of the peHaPi29 ssRNA phage RNA folding threshold (>51.4%) to all 817,234 GenBank viral sequences ≥1,000 nucleotides yielded 56,047 qualifying sequences (6.9%) (Supplemental Table ST4). Ranking these accessions by peHaPi29 score revealed that several non-phage virus groups were overrepresented among the highest-scoring sequences. Most notably, every capsidless ssRNA ormycovirus (266 sequences) exceeded the peHaPi29 threshold, as did 1,528 of the 2,117 narna(-like) viruses (72.2%) (Supplemental Table ST4). Whereas narna(-like) viruses exhibited a broad range of peHaPi29 scores, encompassing both high and low values, all ormycoviruses clustered at the high end of the score distribution (Figure 2A), albeit with substantial overlap with ssRNA phages. Additional folding analyses of these capsidless viruses indicated that the predicted hairpins (Figure 1C–F) are substantially smaller and more uniform in size than those observed in ssRNA phages (Figure 1A and 1B). To better differentiate the newly observed RNA folding structure in capsidless viruses from that of ssRNA phages, we conducted a second peHaPi parameter sweep, contrasting all 266 ssRNA ormycovirus GenBank accessions with 6,336 *Leviviricetes* (ssRNA phages) accessions. A maximum hairpin length of 12 nucleotides (peHaPi12) produced the best discrimination between the two groups with an ROC-AUC of 0.986, an operating threshold of 57.3%, sensitivity of 0.959, and specificity of 0.942 (Supplemental Table ST3). These findings indicate that the pervasive RNA folding observed in ssRNA phages and in ormycoviruses can originate from distinct structural architectures. This is illustrated by the peHaPi12 density plot, in which ormycovirus sequences form a clearly separate cluster from all other sequences (Figure 2B). Notably, ssRNA phage sequences are more strongly separated from ormycoviruses when assessed using peHaPi12 scores (Figure 2B) than with peHaPi29 scores (Figure 2A), highlighting differences captured by the two metrics.

In addition to spanning nearly the entire genome, indicated by the low fraction of unpaired nucleotides (Figure 2E) and the high number of hairpins per 100 nt (Figure 2F), ormycovirus hairpins show notably little variation in stem length within each individual genome sequence as illustrated in Figure 1A–B (phages) and Figure 1C (ormycovirus) (see also Supplemental Figure SF2). Across ormycovirus genomes, the average hairpin stem length spans 7.0 to 15.8 nt (Figure 2C; Supplemental Table ST2). To visualize this pronounced structural uniformity, we plotted each sequence’s peHaPi12 score against the corresponding standard deviation of hairpin lengths (Figure 2D). Treating ssRNA phage sequences as a reference contrast to the ormycovirus dataset, we identified the hairpin-length standard deviation that best balances sensitivity and specificity (TPR vs. FPR), resulting in an optimal threshold of 13.94 nucleotides (Figure 2D).

### Identification of viruses exhibiting the novel pervasive RNA hairpin structural type

Applying the established hairpin folding criteria (peHaPi12 > 0.573 and hairpin-length standard deviation < 13.94 nt) to RNA folds computed for all 817,234 GenBank viral sequences, we identified 1,802 sequences exhibiting RNA-folding characteristics similar to those of ormycoviruses; hereafter termed peHaPi(+) (Supplemental Table ST5; Supplemental Figure SF3). Of these sequences, 665 (37%) correspond to narna(-like) viruses, 267 (15%) to ormycoviruses, and 97 (5%) to ssRNA phages, while the remaining 773 accessions (43%) have inconsistent and/or incomplete annotations. Using a standard BLASTN search against the GenBank core nucleotide (nt) database (Supplemental Table ST5), 83.4% of the selected peHaPi(+) sequences could be assigned to 12 virus classes, leaving 297 accessions with alternative annotations (Table 1). Among the 0.2% of GenBank virus accessions that are peHaPi(+), we observed strong enrichment in ormycoviruses (95%), moderate enrichment in narna(-like) viruses (31%), and near absence in ssRNA phages (1.5%) (Table 1). This pattern is consistent with the thresholds defined to discriminate between ormycovirus and ssRNA phage folding structures (Figure 2D). Nine additional virus classes also contained peHaPi(+) sequences, with frequencies ranging from 2% to 88% within each class. By contrast, the only two other capsidless virus types, mito(-like) virus and botourmia(-like) virus, showed low peHaPi(+) prevalence in their class of 2.7 and 6.1%, respectively. With respect to genome type, (+)ssRNA viruses were predominant overall (69%); however, peHaPi(+) structures were also detected in 30 double-stranded RNA viruses and 229 DNA viruses (Table 1). Notably, of the four DNA virus classes containing peHaPi(+) members, three are pararetroviruses that replicate via an RNA intermediate, whereas the polar freshwater circular DNA virus class is defined by sampling location and provides no information on viral type (Table 1).

**Table 1.**
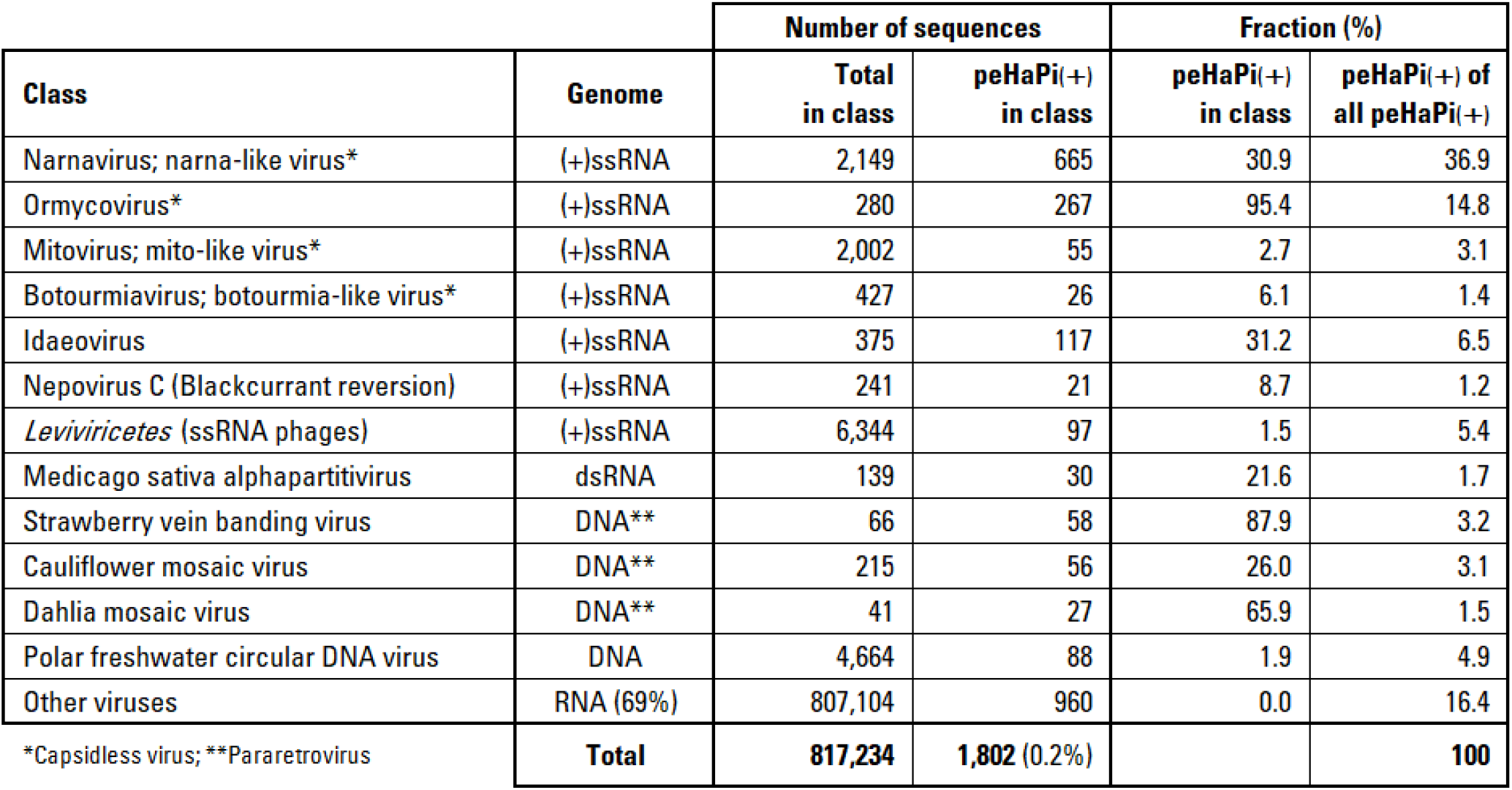
Classification of GenBank virus sequences (>1,000 nt) with pervasive occurrence of hairpins.

| Class | Genome | Number of sequences |  | Fraction (%) |  |
| --- | --- | --- | --- | --- | --- |
|  |  | Total in class | peHaPi(+) in class | peHaPi(+) in class | peHaPi(+) of all peHaPi(+) |
| Narnavirus; narna-like virus* | (+)ssRNA | 2,149 | 665 | 30.9 | 36.9 |
| Ormycovirus* | (+)ssRNA | 280 | 267 | 95.4 | 14.8 |
| Mitovirus; mito-like virus* | (+)ssRNA | 2,002 | 55 | 2.7 | 3.1 |
| Botourmiavirus; botourmia-like virus* | (+)ssRNA | 427 | 26 | 6.1 | 1.4 |
| Idaeovirus | (+)ssRNA | 375 | 117 | 31.2 | 6.5 |
| Nepovirus C (Blackcurrant reversion) | (+)ssRNA | 241 | 21 | 8.7 | 1.2 |
| *Leviviricetes* (ssRNA phages) | (+)ssRNA | 6,344 | 97 | 1.5 | 5.4 |
| Medicago sativa alphapartitivirus | dsRNA | 139 | 30 | 21.6 | 1.7 |
| Strawberry vein banding virus | DNA** | 66 | 58 | 87.9 | 3.2 |
| Cauliflower mosaic virus | DNA** | 215 | 56 | 26.0 | 3.1 |
| Dahlia mosaic virus | DNA** | 41 | 27 | 65.9 | 1.5 |
| Polar freshwater circular DNA virus | DNA | 4,664 | 88 | 1.9 | 4.9 |
| Other viruses | RNA (69%) | 807,104 | 960 | 0.0 | 16.4 |
| *Capsidless virus; **Pararetrovirus | **Total** | **817,234** | **1,802 (0.2%)** |  | **100** |

### Predicted RNA structure stability and ensemble diversity across peHaPi12 scores

The peHaPi scores that were used in the preceding analyses were calculated from the predicted RNA secondary structure with the lowest calculated Gibbs free energy among all structures considered by the RNAfold model [45]. These so-called minimum free energy (MFE) structures do not necessarily represent the exact structures that the virus genomes will adopt under biologically relevant circumstances. Since RNAfold can also calculate a full thermodynamic ensemble of alternative structures, we examined whether increasing peHaPi12 was associated with greater predicted structural stability and reduced ensemble diversity.

A general trend toward more negative MFE per nucleotide was observed with increasing peHaPi12, although this pattern was primarily evident at peHaPi12 values below 0.4 in ssRNA phages and above 0.6 in narnaviruses (Figure 4A). In addition, peHaPi(−) genomes (peHaPi12 < 0.573) exhibited greater variability in MFE than peHaPi(+) genomes, indicating that low-MFE structures can occur in some peHaPi(−) genomes but are more prevalent among peHaPi(+) narnavirus genomes. These subgroup-specific patterns indicate that genomes with higher peHaPi12 scores may, under these conditions, have predicted minimum-energy structures that are modestly more thermodynamically favorable than those of genomes with lower scores. This observation is consistent with work by Terai *et al.* [56], who developed CDSfold to optimize coding sequences for low MFE. Their example of a highly stabilized sequence formed a secondary structure dominated by consecutive short hairpins, qualitatively resembling the peHaPi(+) structural phenotype shown in Figure 1C–F. Nevertheless, these analyses are descriptive, and MFE predictions alone do not establish differences in in vivo RNA folding or stability, nor a causal relationship between thermodynamic stability and peHaPi12.

Lower ensemble diversity suggests that RNAfold predicts a relatively narrow range of similar, thermodynamically probable structures. Higher values suggest a broader structural ensemble in which markedly different secondary structures may be similarly probable. In ssRNA phages, ensemble diversity showed an overall increasing trend with peHaPi12 (Figure 4B). In contrast, above the peHapi(+) classification threshold, narnaviruses showed a reduction in median ensemble diversity with lower variation. Thus, the predicted folding landscape of peHaPi(+) narnaviruses is more strongly concentrated around a single structure or a small family of closely related structures. Collectively, these results indicate that high-peHaPi12 genomes may combine relatively stable predicted secondary structures with more constrained and consistently predicted conformations, particularly in narnaviruses.

### High peHaPi12 scores are associated with altered codon pairing phases

For a peHaPi(+) RNA secondary structure to evolve, the extensive formation of short hairpins must remain compatible with the constraints imposed by protein-coding sequences. To investigate whether particular codon pairing arrangements facilitate this compatibility, we examined the three codon base-pairing phases described by Shabalina *et al.* [58], which capture the possible pairing configurations between codon positions 1, 2, and 3. Previous analysis by Fricke *et al.* [57] showed that phase *c1* is generally under-represented in protein-coding sequences, including viruses, whereas phases *c2* and *c3* occur at similar frequencies. The peHaPi(+) phenotype is characterized by numerous consecutive short hairpins, resulting in base-pairing between codons that are relatively close in the coding sequence. Such structures may preferentially favor the phase *c1* and *c2* pairing arrangements that make optimal use of the more sequence-flexible third, or wobble, codon position. In contrast, phase *c3* pairs the third positions of opposing codons with each other, leaving fewer opportunities for other codon positions to pair with a wobble position. We therefore hypothesized that phase *c3* would become less frequent as the peHaPi phenotype becomes more pronounced.

To test this, we calculated the frequencies of all three codon pairing phases for narnavirus and ormycovirus coding sequences using GenBank annotations and compared them with their peHaPi12 scores (Figure 3C). In these viruses, *c3* pairing was generally less frequent than *c1* and *c2* pairing, contrasting with the previously reported under-representation of *c1* in viruses overall [57]. Moreover, as predicted, above the peHaPi(+) classification threshold of 0.573, the fraction of base pairs belonging to phase *c3* gradually decreased with increasing peHaPi12 score. Although this trend was modest, the progressive reduction in *c3* pairing is consistent with increasing constraints associated with pervasive hairpin formation. Together with the peHaPi12-associated decreases in minimum free energy and ensemble diversity, these results further support the biological relevance of the predicted secondary structures of peHaPi(+) narnavirus and ormycovirus genomes.

**Figure 3.**
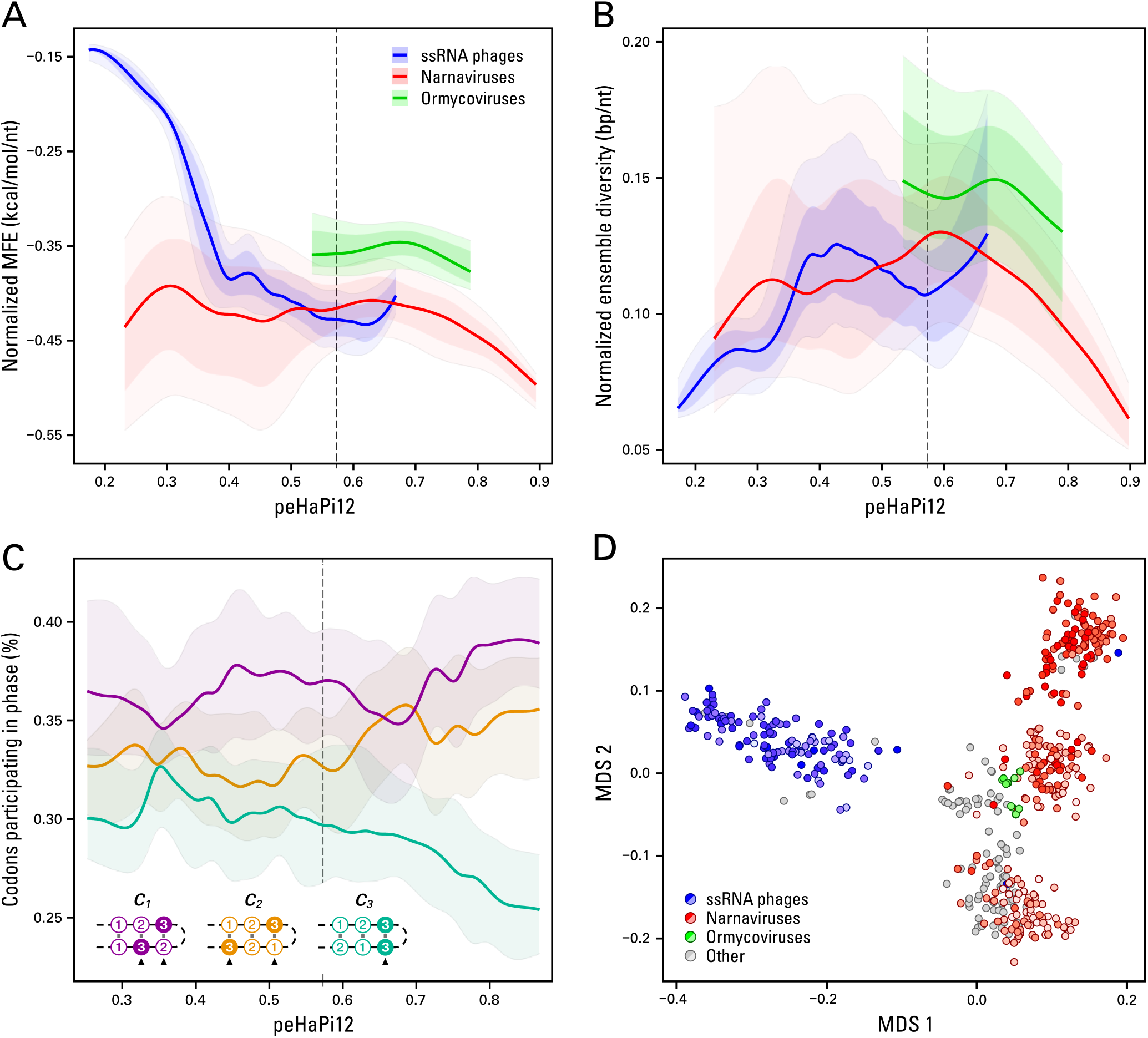
Relationship between peHaPi12 score, RNA secondary-structure and codon base-pairing phase metrics, and RdRp palmprint similarity. Quantile ribbon plots showing the relationship between peHaPi12 score and two RNAfold-derived metrics normalized by genome length: (A) minimum free energy (MFE) per nucleotide and (B) ensemble diversity per nucleotide. The dataset included all GenBank viral sequences ≥1,000 nucleotides annotated as ssRNA phages (blue), narnaviruses (red), or ormycoviruses (green). Within each viral class, sequences were ordered by peHaPi12 score and divided into equal-frequency bins of 10. Solid lines indicate bin-wise medians. Dark and light ribbons represent the 25th–75th and 10th–90th percentile ranges, respectively. Bin-wise summary statistics were interpolated using smoothing splines to visualize trends across peHaPi12 scores. The vertical dashed line marks the peHaPi(+) classification threshold of 0.573. C) Quantile ribbon plots of codon phase versus peHaPi12 score for codons participating in base pairing. For each narnavirus and ormycovirus genome, codons with one or more nucleotides paired to nucleotides in other codons were classified as *c1* (1:1, 2:3, 3:2; purple), *c2* (1:3, 2:2, 3:1; orange), or *c3* (1:2, 2:1, 3:3; teal) following Fricke *et al.* [57], and expressed as a proportion of all codons in that genome. A codon could contribute to more than one phase class if different nucleotides participated in different pairing phases. Viruses were ordered by peHaPi12 score and divided into equal-frequency bins of 20. Solid lines indicate bin-wise medians, and ribbons denote the 25th–75th percentile range. Bin-wise summary statistics were interpolated using smoothing splines. Drawings illustrate the three nucleotide-pairing phases in a schematic stem-loop. Base pairs involving the degenerate third (“wobble”) codon position are marked by black triangles, showing that phases *c1* and *c2* make the most effective use of genetic-code degeneracy. D) Relationship between peHaPi score and RdRp palmprint similarity. Narnaviruses (red), ormycoviruses (green), ssRNA phages (blue), and “Other” viruses (grey) were clustered by amino acid sequence similarity within the RdRp palmprint region [48]. One representative genome, together with its peHaPi12 score, was selected from each cluster, reducing the dataset from 817,234 viral sequences to 4,979 cluster representatives. To reduce overrepresentation of peHaPi(−) sequences, peHaPi(−) narnavirus and ssRNA-phage cluster representatives were randomly subsampled to match the numbers of peHaPi(+) representatives within the corresponding virus class, yielding 537 clusters. Representative palmprint amino acid sequences were aligned, pairwise distances derived from the multiple sequence alignment were calculated, and the resulting distance matrix was projected by multidimensional scaling (MDS). Each point represents the RdRp palmprint sequence of one cluster representative. Viruses assigned to the “Other” category were included only when classified as peHaPi(+). Within each viral-class color, intensity reflects peHaPi12 score, with lighter shading indicating lower scores and brighter shading indicating higher scores.

### Association between RdRp palmprint sequence similarity and peHaPi12 score

We next investigated whether virus genomes with high peHaPi12 scores showed greater sequence similarity to each other than to genomes with low scores. To enable robust comparisons across highly divergent virus classes, we generated a multiple sequence alignment of the RdRp palmprint regions from all narnaviruses, ormycoviruses, ssRNA phages, and peHaPi(+) “Other” viruses. To make the analysis computationally feasible, closely related virus genomes were first grouped into clusters according to amino acid sequence similarity within the RdRp palmprint region, and one representative sequence was selected from each cluster.

The ssRNA phage representatives formed a relatively compact group that was clearly distinct from the other virus categories (Figure 3). Within this group, the distribution of representatives was largely independent of peHaPi12 score, and the two score classes did not form distinct clusters. Thus, the peHaPi phenotype does not appear to be strongly linked to any sequence-defined lineages represented among the ssRNA phages. By contrast, the narnavirus representatives occupied a much broader region of the plot. High-and low peHaPi12-scoring viruses were largely separated, indicating that RdRp palmprint sequence similarity is associated with the peHaPi property within narnaviruses. A distinct intermediate group of narnaviruses contained a mixture of representatives with high and low peHaPi12 scores. The ormycovirus clusters themselves were restricted to a comparatively compact region and were uniformly peHaPi(+), consistent with conservation of extensive genome folding across the sampled ormycoviruses.

The observation that ssRNA phages remained relatively similar regardless of their peHaPi12 scores, whereas peHaPi(+) and peHaPi(‒) narnaviruses were clearly separated, suggests that the peHaPi property may have emerged independently through different evolutionary routes in these virus groups. The marked separation between peHaPi(+) narnaviruses and peHaPi(+) ssRNA phages may be compatible with, but does not by itself demonstrate, convergent evolution of the peHaPi property in *Leviviricetes* and non-*Leviviricetes*.

### Possible functions of the pervasive hairpin structure

It is clear that pervasive hairpin structures exist in various viral lineages (Figure 1 and 2). In many peHaPi(+) sequences, arrays of short hairpins tile almost the entire genome, often separated by at most two nucleotides. Achieving such a densely packed, repetitive RNA architecture throughout a virus genome is nontrivial, given that the successive palindromic sequences that make up the stem of each hairpin, most of the time still need to contribute to a functional open reading frame (ORF). Moreover, many narnavirus genomes show a striking uniformity in hairpin length with stem lengths ranging from seven to 14 nucleotides (Figure 1E, 1F, 2E, 2F, and Supplemental Figure SF4). These observations strongly suggest selection for a specific, length-dependent and hairpin-spacing dependent role rather than incidental RNA folding.

#### Hairpin-loop sequence analysis

There are several conceivable functions for the peHaPi(+) structures. A first possibility is that the hairpins act as a structural scaffold for coat proteins during virion construction. This model is well established for ssRNA phages [11, 25–28], and as shown, these viruses have a peHaPi structure that is distinct from other virus sequences. By analogy, one might expect a similar scaffolding function for peHaPi(+) structures in the viruses analyzed here. However, because over half of the peHaPi(+) viruses we identified are capsidless viruses, this possibility seems less likely, unless these viruses hijack capsids from co-infecting viruses or instead rely on proteins of the host organism to form protective ribonucleoprotein (RNP) complexes.

We evaluated the RNP scaffolding model by comparing hairpin loop sequences from peHaPi(+) narna(-like) viruses with those identified in peHaPi(‒) narna(-like) viruses. The rationale was that, if numerous copies of the same protein specifically bind these hairpins, their recognition might depend on a conserved sequence motif within the loops. An example for such RNA-protein interactions is found in bacteriophage MS2, whose RNA genome contains more than 50 stem-loops, most of which contact the capsid through their loop regions [59]. However, analysis of single nucleotide, triplet, and tetramer frequencies within the loop regions provided no convincing evidence for a conserved, sequence-specific docking motif in the hairpin loops of peHaPi(+) viruses (Supplemental Table ST6).

#### vsiRNA response to peHaPi(+) viruses

A plausible function of peHaPi folding is protection of the viral genome from the host antiviral siRNA response. The dense arrangement of short hairpins could interfere with antiviral RNA interference at two distinct stages. First, extensive intramolecular folding may reduce the formation of long viral double-stranded RNA intermediates, which are typically required for Dicer-dependent production of virus-derived siRNAs (vsiRNA) [60, 61]. Although the viral RNA polymerase must temporarily unfold local hairpins while synthesizing the complementary strand, both RNA strands are expected to rapidly refold into the characteristic peHaPi structure after emerging from the polymerase [62]. This rapid refolding would limit their capacity to re-anneal into extended, Dicer-accessible duplexes. Second, peHaPi folding may reduce the accessibility of the viral genome to vsiRNA-loaded RISC complexes. Efficient Argonaute-mediated targeting requires an exposed sequence that permits initial pairing with the approximately seven-to eight-nucleotide seed region, followed by extension of guide-target complementarity. Stable local secondary structures can inhibit this process by creating barriers to duplex formation and RISC-mediated cleavage [63–65]. In principle, single-stranded siRNAs derived from peHaPi(+) RNA might also fold back on themselves, reducing their antiviral activity. However, functionally active siRNAs are largely maintained in a defined, extended conformation within Argonaute [66], making siRNA self-folding an unlikely major contributor to immune evasion. Overall, peHaPi folding may suppress both vsiRNA biogenesis and subsequent recognition of the viral genome.

To assess whether peHaPi folding completely prevents the induction of an antiviral siRNA response, we revisited one of our earlier studies in which we identified the novel ormyco-like virus Alaria esculenta virus NL-1 in an *Alaria esculenta* seaweed sample [67] (Supplemental Table ST5). Both viral segments exhibit a peHaPi(+) architecture, with peHaPi12 scores of 0.770 for genome segment RNA1 (PP793779) and 0.766 for RNA2 (PP793780). Nearly all smallRNA-seq reads aligning to RNA1 and RNA2 were 21 nt in length and carried a 5′-terminal T nucleotide [67], two characteristic features of virus-derived siRNAs. This strong enrichment indicates that the host generated a pronounced antiviral siRNA response, providing evidence that peHaPi(+) viruses can trigger RNAi-mediated immunity in seaweed.

To determine whether other peHaPi(+) viruses are capable of eliciting an siRNA response, we generated smallRNA-seq, RNA-seq, and DNA-seq datasets from 48 plant samples collected from a Dutch botanic garden. Its tropical climate was expected to support a broad diversity of fungi, which represent the principal hosts of narna(-like) and ormyco(-like) viruses. RNA-seq and smallRNA-seq reads were mapped against the 1,802 peHaPi(+) accessions retrieved from GenBank. Among these accessions, 50 displayed appreciable smallRNA-seq read mapping indicative of siRNAs, whereas 102 showed RNA-seq read mapping consistent with viral RNA (vRNA) presence (Supplemental Table ST7). Integration of both datasets identified only eight peHaPi(+) GenBank accessions for which both viral presence and corresponding siRNAs were detected. However, even among these accessions, the maximum siRNA mapping coverage extended to only slightly more than 28% of the complete accession sequence (Supplemental Table ST7). These limited and fragmentary mapping patterns indicate that most peHaPi(+) GenBank viruses were absent from the analyzed samples, while the few viruses that may have been present yielded insufficient read numbers for a reliable assessment. Consequently, this dataset did not permit any conclusion regarding the siRNA response elicited by peHaPi(+) viruses.

As a complementary approach, we subsequently performed *de novo* discovery of peHaPi(+) viruses within the botanic garden experiment. Of 669 distinct viral contig sequences identified (<u>Supplemental Document SD2</u>), 74 were classified as peHaPi(+). Among these, 34 (46%) displayed substantial siRNA representation, with sequence coverage ranging from 30 to 100% (Supplemental Table ST8; Supplemental Figure SF5). Comparable evidence of siRNA involvement was observed for 192 of the 595 (32%) peHaPi(‒) viral contigs. Thus, the widespread and occasionally abundant detection of siRNAs derived from peHaPi(+) viruses (Supplemental Table ST8) argues against the hypothesis that the peHaPi structure markedly modifies the siRNA response directed against these viruses.

### Using the peHaPi structure to identify new virus sequences

The botanic garden experiment supports the use of the peHaPi(+) scoring criteria as a means of detecting candidate viral contigs that may not be recognized by conventional sequence homology-based approaches. Among the 90 peHaPi(+) contigs, protein-level similarity to narnaviruses and ormycoviruses was detected for 44 and 18 contigs, respectively (Table 2; Supplemental Table ST8). In contrast, only 27 of the 595 peHaPi(‒) viral contigs showed protein similarity to narnaviruses, while only 1 showed similarity to ormycoviruses. These relative frequencies were consistent with the representation of these virus groups in GenBank (Table 1). Of the remaining peHaPi(+) contigs, 12 showed similarity to other virus types, whereas 16 had no detectable similarity to known viruses and therefore represent candidates for highly divergent, previously undescribed narna(-like) or ormyco(-like) viruses (Supplemental Table ST8).

**Table 2.**
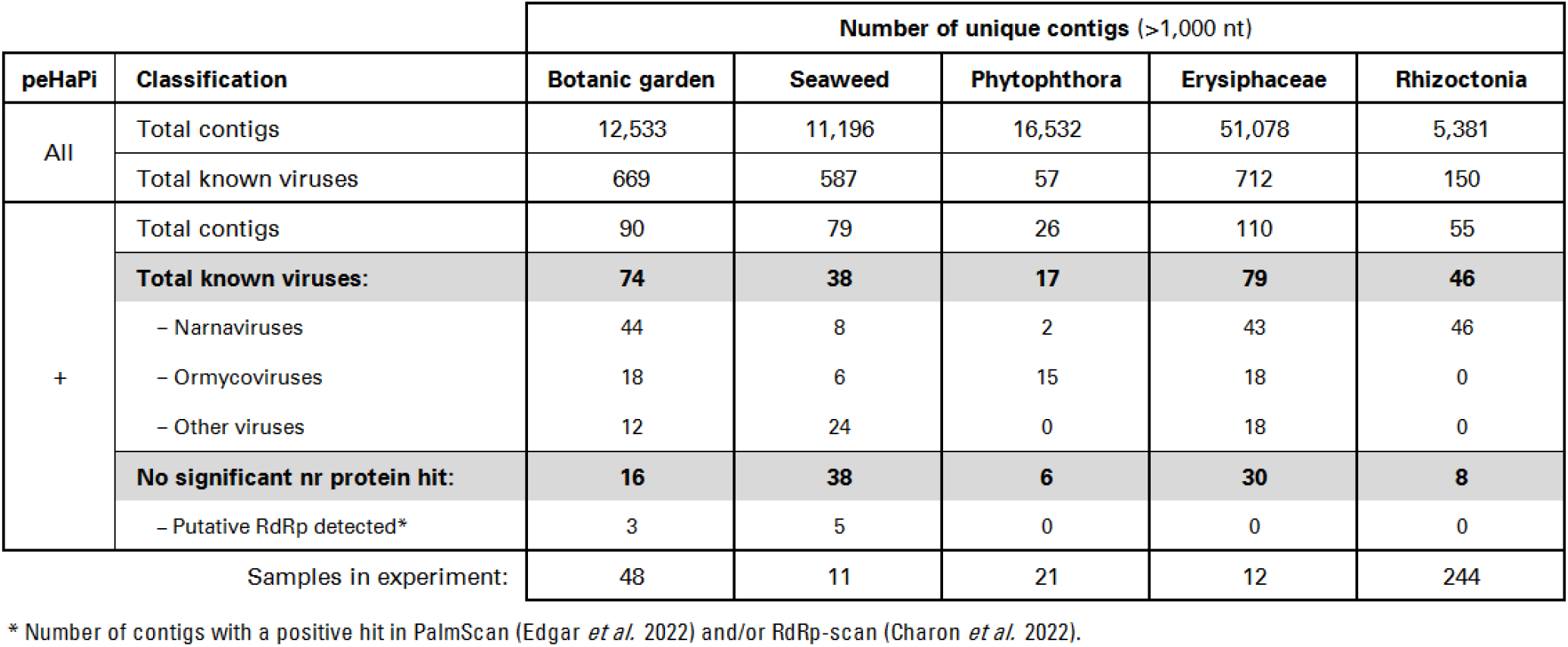
Identification of peHaPi(+) viral contigs across independent RNA sequencing datasets.

The large number of distinct contigs generated by RNA-seq makes exhaustive RNAfold-based secondary-structure prediction computationally demanding. To enrich for RNA contigs of viral origin, we routinely sequence DNA alongside RNA and exclude contigs with detectable mapping of genomic DNA reads, hereafter referred to as DNA(+) contigs [68, 69]. This removes transcripts derived from DNA genomes present in the sample, including the host genome and other DNA-based organisms, thereby enriching for DNA(−) contigs more likely to originate from RNA viruses. Across the 48 botanic garden samples, this filtering reduced the number of contigs requiring further analysis from 959,802 to 12,533 DNA(−) candidates. Combining DNA(−) filtering with peHaPi(+) classification could therefore provide a highly selective strategy for identifying divergent narna- and ormycovirus candidates. For this approach to be robust, however, host-derived transcripts should only rarely satisfy the peHaPi(+) criteria. The excluded DNA(+) fraction provides a large set of predominantly host-derived transcripts for estimating this false-positive rate. Applying the peHaPi(+) criteria to the 947,269 DNA(+) contigs identified only 36 peHaPi(+) contigs (0.004%; ∼1 in 26,000) (Supplemental Table ST9; <u>Supplemental Document SD3</u>; and Supplementary Figure SF6). Two of these, including the contig with the highest peHaPi score, showed similarity to ormycoviruses, whereas the remainder appeared to originate from host plant genes encoding hypothetical proteins. Thus, host-derived transcripts only rarely satisfy the peHaPi(+) criteria, supporting the combined use of DNA(−) filtering and peHaPi(+) classification for selective identification of candidate RNA virus sequences.

We next evaluated the combined DNA(−)/peHaPi(+) strategy using four additional independent seaweed [37] and fungal [70–72] datasets (Table 2 and Supplemental Table ST1). Two datasets, comprising various *Rhizoctonia* [71] and Erysiphaceae [70] samples, had previously been examined in separate studies that identified numerous new narna- and ormycoviruses. Because these datasets were already known to be rich in these virus groups, they provided a suitable test of whether the new strategy could reveal additional candidates missed by earlier analyses. Across all datasets, including the botanic garden data, 17 to 79 peHaPi(+) contigs per dataset showed similarity to known narna- and ormycoviruses or other viruses (Table 2). In addition, 98 peHaPi(+) contigs lacked detectable protein homology to known viruses, corresponding to 6 to 38 candidates per dataset. Only 8 of these 98 contigs contained an RdRp signature detectable by either PalmScan [48] or RdRp-scan [73]. The remaining 90 contigs would therefore have escaped identification by established protein homology and RdRp detection approaches.

The consistent recovery of homology-negative candidates across biologically distinct, independently generated datasets supports the broader applicability of the peHaPi metric for identifying highly divergent RNA virus sequences that may escape conventional sequence homology-based detection.

## Concluding remarks

In this study, we discovered a remarkable RNA structure that occurs predominantly in capsidless viruses. This RNA structure, which we quantify using our newly developed peHaPi metric, is characterized by a high density of relatively short hairpins of uniform length that span the entire viral genome. Across peHaPi(+) viruses, mean hairpin stem length varies from approximately 7 to 16 base pairs, whereas stem lengths within individual genomes are remarkably uniform. This combination of strong within-genome conservation and substantial between-virus variation suggests that each virus may favor a characteristic stem length while independently modulating hairpin abundance and maintaining a functional RdRp ORF.

Sequence homology-based virus discovery is inherently limited for highly divergent viruses [74, 75]. peHaPi provides a complementary, structure-based approach that enables homology-independent identification of ormycoviruses and a subset of narnaviruses. Recent studies have similarly exploited RNA structural information for RNA virus discovery, using ribozyme covariance models, conserved local stem-loops inferred from sequence variation, or genome-scale thermodynamic stability [76–78]. In contrast, peHaPi directly quantifies a specific genome-wide architecture of short, uniformly sized hairpins in individual sequences, enabling selection of capsidless RNA-virus candidates without requiring a known structural motif, cross-sequence covariation, or detectable viral protein homology. However, the peHaPi scoring framework could be further refined to improve discrimination, for example by incorporating potential pseudoknot formation, which is not captured by RNAfold. In addition, peHaPi organization forms a continuum from moderate to highly pronounced dense-hairpin patterns rather than a naturally discrete classification. An absolute peHaPi(+)/peHaPi(−) boundary is therefore not evident, and the current threshold was empirically defined based on confirmed ormyco- and narnaviruses.

Although peHaPi(+) organization occurs predominantly in capsidless RNA viruses, we also identified DNA viruses that exhibit a clear peHaPi(+) pattern (Table 1), indicating that this structural principle is not restricted to capsidless RNA viruses. The presence of peHaPi(+) structures in both dsRNA and DNA viruses points to a selective pressure that extends beyond the genomic RNA itself and likely acts on viral mRNAs as well. This idea is further supported by the observation that several of the DNA viruses are pararetroviruses that rely on an RNA intermediate in their replication cycle [79]. Thus, selection may act at multiple stages of the viral life cycle, shaping RNA secondary structure for optimal function in replication, translation, or genome stability. The coordinated regulation of hairpin abundance and structural dimensions further suggests that both properties are subject to evolutionary selection and may contribute to as-yet-unidentified biological functions.

Protection against the host siRNA-mediated antiviral response appears at first sight to be an attractive explanation for the evolution of the peHaPi architecture. However, in the large botanic garden experiment we did not observe a reduced siRNA response for peHaPi(+) viruses compared to peHaPi(−) viruses. This lack of differential targeting suggests that siRNA evasion alone cannot account for the emergence or maintenance of the peHaPi structure, and that other functional roles must be considered.

How the extensive arrays of highly uniform secondary structures in peHaPi(+) genomes organize into a defined tertiary structure remains unresolved. Although experimental structures containing several adjacent stem-loops are available, we are unaware of any that reproduce the long, closely spaced series of similarly sized hairpins characteristic of peHaPi(+) genomes. Examples include three consecutive stem-loops in the Moloney murine leukemia virus packaging signal, six stem-loops in RsmZ, and numerous stem-loop or helical elements resolved within MS2 and satellite tobacco mosaic virus genomes [59, 80–82]. However, these structures either contain only a few adjacent hairpins, are embedded in protein-bound complexes, or lack sufficient RNA connectivity to reveal how a long series of hairpins collectively organizes an RNA molecule.

Consequently, it remains unclear whether the consecutive hairpins in peHaPi(+) genomes behave largely as independent local structures or interact through loop-loop contacts, coaxial stacking, or other tertiary interactions to compact the RNA into a defined architecture. Such an organization could potentially compensate for the lack of a protective capsid while not being subject to the packing constraints imposed by a capsid [35, 61]. Tertiary interactions are well established as organizers and regulators of RNA structure, and long-range intragenomic RNA-RNA interactions can control translation, replication and transcription in RNA viruses [83]. We therefore speculate that the pervasive peHaPi organization may ultimately function by imposing a characteristic tertiary architecture on the viral genome, potentially influencing RNA stability and intracellular maintenance, accessibility to host or viral proteins, replication-complex assembly, translation, or recognition by host defense systems. Experimental determination of this architecture will be required to test these possibilities. Future approaches combining nucleotide-resolution structure probing, mapping of long-range RNA-RNA interactions, structural biology and experimentally constrained 3D RNA modelling [84] should help establish whether high peHaPi scores correspond to a distinctive genome-scale tertiary fold and clarify its contribution to the biology and evolution of these predominantly capsidless RNA viruses.

## Author contributions

T.M.B., R.J.D.: Conceptualization, Writing – Original Draft, Visualization, Supervision. R.J.D., M.F.v.O., W.A.E., S.M.v.L., M.J.J.: Investigation. W.C.d.L., R.J.D., M.J.J.: Software, Formal analysis, Visualization. R.J.D., M.J.J.: Writing – Review & editing.

## Acknowledgements

We thank prof. dr. Klaas Timmermans and Job Cohen (Department of Estuarine and Delta Systems, NIOZ Royal Netherlands Institute for Sea Research, Yerseke, the Netherlands) for providing seaweed samples, and dr. Roderick Bouman and Rogier van Vugt (Hortus botanicus Leiden, Leiden University, Leiden, the Netherlands) for their assistance with botanical garden sample collection and annotation.

## Supplemental information

All supplemental files are available through FigShare (https://figshare.com/s/749c39c3008c8666cbc4).

### Supplemental Documents

**SD1.** Custom R scripts

**SD2.** Hortus contigs DNA(-)

**SD3.** 36 sequences of DNA(+) and peHaPi(+) contigs Hortus RNA-seq

### Supplemental Figures

**SF1.** peHaPi rules and example calculation

**SF2.** Hairpin length vs peHaPi score

**SF3.** 1802 accessions RNAfold

**SF4.** 10 folding examples with different stem length

**SF5.** 133 contigs Hortus peHaPi(+) RNAfold

**SF6.** 36 contigs Hortus DNA(+) peHaPi(+) RNAfold

### Supplemental Tables

**ST1**. Description sequencing data and reference genomes

**ST2**. 817,235 GenBank virus sequences > 1,000 nt

**ST3**. peHaPi parameter sweeps to discern different types of pervasive hairpin RNA structures

**ST4**. 5,607 GenBank virus sequences with a peHaPi29 score higher than 51.4%

**ST5**. 1,802 GenBank virus sequences with a similar RNA folding structure as ormyco(-like) viruses

**ST6**. Distribution of nucleotides in the hairpin loops of 2,118 narna(-like) virus sequences

**ST7**. Combined mapping of siRNA reads and vRNA-reads to peHaPi(+) GenBank accessions

**ST8**. Mapping siRNA reads to RNA-seq contigs derived from a 48 sample botanic garden experiment and selected for no DNA-seq read mapping

**ST9**. 36 RNA-seq contigs with mapping DNA-seq reads derived from the botanic garden experiment that met the peHaPi(+) criteria.

